# The alterations of the synthetic pathway and metabolic flux of auxin indole-3-acetic acid (IAA) govern thermotolerance in *Lentinula edodes* mycelia subjected to heat stress

**DOI:** 10.1101/2025.04.09.648085

**Authors:** Xiaoxue Wei, Jiaxin Song, Jiayue Chen, Yang Xiao, Yan Zhou, Yinbing Bian, Yuhua Gong

**Author notes:** Address correspondence to Yuhua Gong. Present address: Xiaoxue Wei, Jiaxin Song, Jiayue Chen, Yang Xiao, Yan Zhou, Yinbing Bian, Huazhong Agricultural University, No.1 Shizishan Street, Hongshan District, Wuhan, 430070, Hubei Province, China.

## Abstract

Heat stress poses a significant constraint on the annual production of *Lentinula edodes(L. edodes)*, a challenge that has been intensified by global warming. Previous studies have established a close relationship between intracellular indole-3-acetic acid (IAA) content and heat tolerance in *L. edodes*. However, the specific changes in the IAA synthesis pathway and the target genes regulated by IAA under heat stress remain unclear. We employed targeted metabolomics and transcriptomics analysis to investigate the alterations in IAA synthesis pathways and gene expression in both heat-tolerant and heat-sensitive strains at various time points during heat stress. Our findings revealed that IAA is primarily synthesized via the tryptamine (TAM) and indole-3-pyruvic acid (IPYA) pathways in *L. edodes*. Heat-sensitive strain Y3357 exhibited excessive accumulation of tryptamine after heat stress. Silencing of the key genes of IAA synthesis include tryptophan decarboxylase (TDC) and tryptophan transaminase (TAA) in strain S606 could reduce the thermotolerance of *L. edodes* mycelia. Transcriptome analysis revealed that heat-tolerant strain S606 had an earlier response to protein folding and mitochondrial gene expression compared to the heat-sensitive strain Y3357. Additionally, most genes in the MAPK signaling pathway were up-regulated after 10-24 hours of heat stress, with auxin response elements (AREs) identified in their promoters. These results suggest that the excessive tryptamine accumulation is the newly discovered limiting factor for thermotolerance, and the expression levels of key genes in the IAA synthesis pathway could directly influence hyphal thermotolerance. This study provides a new perspective on the mechanism by which IAA and the synthesis precursors affects thermotolerance of *L. edodes*.

**Importance:** As an important plant hormone, the potential role of IAA in enhancing the heat resistance of *L. edodes* strains has garnered significant attention. This study systematically investigated the intracellular IAA biosynthesis pathway and its metabolic flows in *L. edodes* under varying durations of thermal stress, with particular emphasis on temporal gene expression patterns. Research has demonstrated that excessive accumulation of tryptamine may impair the heat stress recovery capability of *L. edodes*. In contrast, IAA can improve its thermotolerance by modulating the expression of genes associated with the MAPK signaling pathway.

## Introduction

Indole-3-acetic acid (IAA), a crucial phytohormone of the auxin class, plays pivotal roles in multiple aspects of plant growth and development. It actively participates in the regulation of embryo development (1), root organogenesis (2), environmental responses (3), adaptive growth (4), as well as cell division (5) and differentiation (6) processes. High temperature-induced heat stress significantly impairs plant growth and development, ultimately leading to substantial reductions in crop yield and productivity.

Thermomorphogenic responses in plants involve coordinated regulation of auxin biosynthesis and transport following heat stress. Research across multiple species including soybean (7), Arabidopsis (8), lettuce (9), and cucumber (10) demonstrates significant auxin accumulation under thermal stress. The central thermomorphogenesis regulator PHYTOCHROME-INTERACTING FACTOR 4 (PIF4) (11) orchestrates this process by upregulating key auxin biosynthesis genes (TAA1 family) while simultaneously modulating polar auxin transport through PIN-FORMED 1 (PIN1) (12), and PIN2 (13) regulation. This molecular mechanism enables plants to optimize growth patterns and developmental plasticity under elevated temperature conditions.

Contemporary studies have demonstrated the regulatory potential of exogenous auxin application in plant physiology and stress responses. Notably, IAA supplementation has been shown to enhance the growth of soybean seedling (7), promote hypocotyl elongation in Arabidopsis (8), and mitigate heat-induced male sterility in wheat through improved grain retention (14), reduce the loss of grain, increase lettuce shoot (9). Furthermore, auxin application exhibits protective effects against thermal stress in rice cultivation, effectively maintaining pollen viability, spike fertility, and yield components under elevated temperatures (15).

Microbial-derived IAA has been documented to enhance plant resilience to various abiotic stresses including saline, heavy metal (16), and drought stress (17). Recent investigations into thermal stress responses reveal that IAA production serves as a key adaptation mechanism in microorganisms such as *Aspergillus japonicus EuR-26* (18), *L. edodes* (19).

Specifically, our previous research identified a significant increase in endogenous IAA levels in *L. edodes* following 24-hour exposure to 40 °C heat stress. Subsequent experiments demonstrated that exogenous application of IAA and its analogs 0.01mmol/L 2,4-dichlorophenoxyacetic acid [2,4-D]) significantly improved thermotolerance in selected *L. edodes* strains under identical thermal conditions (20). The above results indicate that the intracellular IAA content is closely related to hyphal heat tolerance, but the intracellular IAA synthesis pathway and the mechanism of IAA regulation of heat tolerance are still unclear. Current understanding of IAA biosynthesis reveals two primary routes in biological systems: tryptophan-dependent and tryptophan-independent pathways.

The tryptophan-dependent pathway involves L-tryptophan(Trp) conversion through five distinct enzymatic routes (21). The transformation process mainly consists of five pathways, namely IPYA, TAM, IAM (indole-3-acetamide), IAN (indole-3-acetonitrile), and TSO (Tryptophan side-chain oxidase) pathway (19, 22-27). The IPYA pathway, recognized as a principal auxin synthesis mechanism in microorganisms and plants, employs tryptophan aminotransferase (TAA/TARs) (28, 29) and flavin monooxygenase (YUCCAs) (30) to convert Trp to IPYA (31), subsequently transformed to IAA or via indole-3-acetaldehyde intermediates through pyruvate decarboxylase (IPDC) and indole-3-acetaldehyde (IAALD) dehydrogenase (ALDH) enzymes, as observed in *Ustilago maydis* (32) and *Tricholoma vaccinum* - *Spruce Ecto-mycorrhiza* (33). The TAM pathway initiates with tryptophan decarboxylase (TDC) (34) mediated formation of tryptamine, sequentially converted through N-hydroxy-tryptamine (N-TAM), indole-3-acetaldoxime (IAAOx), and IAALD (indole-3-acetaldehyde) to final IAA production (35), with functional validation in *Taphrina deformans* (36). The IAM pathway utilizes the IaaM (encoding tryptophan monooxygenase) and IaaH (encoding indole-3-acetamide hydrolase) (37) to convert tryptophan to IAM and subsequently to IAA, originally identified in *Pseudomonium* (38)and later confirmed in *Bradyrhizobium japonicum* (39) *Rhizobium* sp. strain NGR234 (40), *Burkholderia pyrrocinia* JK-SH007(41). The IAN pathway, predominantly studied in bacteria, involves the conversion of Trp to IAAOx by cytochrome P450 monooxygenases (CYP79B1, B2, B3) (42, 43) followed by transformation into IAN and subsequent hydrolysis to IAA by nitrilase encoded by the *NIT* (encoding nitrilase) gene. This pathway has been identified in *P. fluorescens* EBC191 and *Alcaligenes faecalis* JM3 (44-46). The TSO pathway has only been shown in *Pseudomonas fluorescens*, and in this pathway Trp is directly converted to IAALD by passing IPYA (25). Comparatively, research on the tryptophan-independent pathway remains limited, though elevated IAA levels detected in Arabidopsis tryptophan biosynthesis mutants suggest the existence of alternative synthesis routes (47).

Given the critical role of auxin in plant growth and development, key genes in its biosynthesis, such as *TDC* and *TAA*. *TDC* gene mediates Trp-to-TAM conversion, with overexpression studies demonstrating enhanced cell elongation in tobacco BY 2 cells (48), improved drought and salt stress tolerance in *Paeonia lactiflora Pall* (49), promoted the root length, root surface area and leaf thickness of tomato (50), delayed senescence and improving resistance to pathogen infection in rice (51).

*TAA* gene catalyzes the critical Trp-to-IPYA conversion in auxin biosynthesis, expression modulated by environmental stimuli such as shade avoidance responses to strong light stress (52). The expression of *TAA* gene is regulated by environmental factors such as temperature signaling (53). Genetic studies in deletion mutants of *TAA* family alleles (*sav3*, *wei8*, *tir2*) (54, 55) reveal its pleiotropic effects on plant development, including altered tiller number but reduction in grain number and size (56), exhibits a defective root gravitropic response (GR) and an increased resistance to CK in primary root growth (57), as well as auxin content (58). Exogenous addition of IAA can complement these phenotypic defects.

Despite these advances in plant and bacterial, significant research gaps persist regarding IAA biosynthetic pathways and their regulatory roles in edible fungi, current limitations include incomplete characterization of fungal-specific IAA synthesis genes, undefined IAA synthesis pathway predominance under thermal stress, and insufficient understanding of cross-talk between auxin signaling and heat response pathways in *L. edodes*. *L. edodes* have emerged as a significant horticultural crop owing to their distinctive flavor and high nutritional value (59), High temperature has a great negative influence on the cultivation of *L. edodes*, causing mycelium damage and stem rot, thus affecting yield, usually resulting in a reduction of 30% (60). In previous studies, shiitake edodes regulated hyphal thermotolerance by regulating intracellular IAA content, but the intracellular IAA synthesis pathway and the mechanism of IAA regulating thermotolerance are unknown. This study analyzed the dynamic changes of plant hormone and gene expression after 40 ℃ heat stress 2, 4, 6, 12, 14 hours by quantitative metabolome and transcriptome methods. This study systematically investigated the intracellular IAA biosynthesis pathway and its metabolic flows in *L. edodes* under varying durations of thermal stress, with particular emphasis on temporal gene expression patterns. Through targeted silencing of two pivotal biosynthetic genes tryptophan aminotransferase (TAA) and tryptophan decarboxylase (TDC). We elucidated their regulatory roles in hyphal thermotolerance. This investigation not only elucidates the mechanistic basis by which auxin modulates thermotolerance in *L. edodes* hyphae under heat stress, but also establishes a novel insight for understanding how auxin homeostasis coordinates heat stress adaptations through metabolic flux regulation and differential gene transcriptional expression within the IAA biosynthesis network.

## Experimental Procedures

### Strains and culture conditions

*L. edodes* YS3357 and S606 strains used in this study were supplied and preserved exclusively by the Institute of Applied Fungi Research, Huazhong Agricultural University. *L. edodes* mycelia were cultivated on MYG medium (containing 20 g malt extract, 20 g glucose,1 g tryptone,1 g yeast extract, 20g agar, 1 L ddH_2_O) in 9 mm petri dish at 25 ℃ for 8 days. Subsequently, mycelia together with petri dish were transferred to the incubator and cultured at 40 ℃ for 0 h, 2 h, 4 h, 6 h, 8 h, 12 h, and 24 h. After 40 ℃ heat stress, *L. edodes* mycelia recovered at room temperature, followed by preservation in liquid nitrogen for subsequent analyses. Simultaneously, fungal mycelia grown at 25 ℃ were subjected to above-mentioned heat stress for RNA extraction and subsequent gene expression analysis (61).

### Quantitative analysis of plant hormones and related metabolites

*L.edodes* strains YS3357 and S606 subjected to 40 ℃ heat stress for different duration were used for metabolite extraction. Extraction and determination of metabolites were performed by Wuhan Metware Biotechnology Co. Ltd. A total of 0.1 g hypha samples were extracted with 1 mL of 70% methanol solution at 4 ℃ for 24 h. The extract was vortexed for 10 min and centrifuged for 5 min (12000 r/min, and 4 ℃), and the supernatant was transferred to clean plastic microtubes, followed by air dryness. The dried samples were dissolved in 100 µL 80% methanol (V/V) and filtered through a 0.22 μm membrane filter for further LC-MS/MS analysis (62). Metabolites were detected using ultra-performance liquid chromatography (UPLC) (Shim-pack1 https://www.metware.cn UFLC SHIMADZU CBM30A) and tandem mass spectrometry (MS/MS) (Applied Biosystems 6500 QTRAP, Wuhan, China). The column temperature was 40 ℃; the flow rate was 0.4 mL·min^−1^; and the injection volume was 2 µL. Mobile phase A was 0.04% acetic acid aqueous solution, and mobile phase B was acetonitrile containing 0.04% acetic acid. The gradient elution was carried out with elution conditions presented in Supplementary Table 1. The voltage of the mass spectrometer was 5,500 V; the temperature of the electrospray ion source was 500 ℃, and the curtain gas was 25 psi. Qualitative analysis of samples was performed according to the secondary spectrum information in the MEDB (metware database). The relative content of metabolites was calculated according to the corrected mass spectrum peak area. The differentially accumulated metabolites (DAMs) between different groups was identified with the thresholds of |log2 FC (fold change)| ≥ 2 and p < 0.05.

Indole-3-acetaldehyde (IAALD), an intermediate metabolite in IAA synthesis, was tested by using a ELISA kit (Shanghai Hengyuan Biotechnology Co., Ltd., Shanghai, China). Briefly, 50 µL samples were added to the standard on the coated plate; 10 µL of the sample dilution (5 x final dilution of the sample) was added to the 96-well plate, added with 50 µL of enzyme labeling reagent (except for blank wells), gently shaken to mix evenly. The microplate was covered with plate sealing film and incubated at 37 ℃ for 60 min. The 30 x concentrated washing solution was 30-fold diluted with distilled water; the plate sealing film was carefully removed; the liquid was discarded; the samples were spin-dried; the washing liquid was added into each well; the samples were stood for 30 s and precipitated, and the supernatant was discarded. The above procedures were repeated 5 times, and then the samples were spin-dried. The dried samples were added with 50 µL color development reagent A, then added with 50 μL of color development reagent B, gently shaken, mixed at 37 ℃, followed by 15-min color development, and finally added with 50 µL of stop solution to terminate the reaction (blue turning yellow). The absorbance (indicated by optical density, OD) at 450 nm was measured within 15 min after addition of stop solution. Afterwards, the standard curve was plotted with the concentration of the standard as the abscissa and the OD value as the ordinate. Then actual sample concentration was calculated according to the OD value and standard curve.

### Total RNA extraction and reverse transcription

The 0.1 g of *L. edodes* fresh hypha was taken, added with liquid nitrogen, quickly ground into powder, transferred to RNase-free 1.5 mL centrifuge tube, added with 600 µL RNAiso Plus, mixed evenly, and stood at room temperature for 5 min, added with 0.2 volume of chloroform, mixed evenly again for 20 s, stood again at room temperature for 5 min, and centrifuged at 13,000 g for 10 min. The supernatant was transferred into a new RNase-free 1.5 mL centrifuge tube, added with 2 / 3 volume of isopropyl alcohol, mixed evenly, stood at room temperature for 10 min, and centrifuged for 10 min at 4 ℃, The speed of 13000 for 10 min. After the supernatant was removed, the precipitate was eluted with 1 mL 70% ethanol twice with the residual ethanol absorbed, air-dried at room temperature on an ultra-clean working table for 5 min, then added with 30 µL RNase-free ddH_2_O, subjected to 65 ℃ water bath, stored at -80 ℃ for subsequent transcriptome sequencing. The sample concentration was determined using an ultra-trace UV spectrophotometer, and the RNA integrity was measured by 1% agarose gel electrophoresis. The reverse transcription of RNA was performed using the HiScript II Q RT SuperMix kit (Vazyme, Nanjing, China) at 42 ℃ for 2 min in 8 µL reaction system containing 400 ng total RNA, 2 µL 4 × gDNA wiper Mix, and RNase-free ddH_2_O. In order to achieve high reverse transcription efficiency, the template was mixed with the RNase-free ddH_2_O. The mixture was homogenized and incubated at 65 ℃ for 5 min, then added with another 4 × gDNA wiper Mix to remove gDNA, added with 2 µL 5 HiScript II qRT SuperMix II, and gently mixed evenly. The subsequent reverse transcription procedures were as follows: at 25 ℃ for 10 min, 55 ℃ for 30 min, and 85 ℃ for 5 s. The reverse transcription product was 20-fold diluted and used for later qRT-PCR analysis.

### Identification of differential expressed genes (DEGs) and cluster analysis

The samples used for transcriptome analysis were collected by the same method as mentioned above for the metabolomics analysis. After RNA extraction, the library was constructed. Sequencing was carried out using the Illumina HiSeq platform. After the quality control of the library, 150 bp paired-end clean reads were obtained. The sequence alignment was conducted with the L808 genome as the reference sequence (63). The genes were counted using Feature Counts software, and the gene expression level was expressed as FPKM (fragments per kilobase of transcript per million mapped reads). The differential expressed genes (DEGs) between comparison groups were identified usingDESeq2 software (64), with the thresholds of |log2 FC (fold change)| ≥ 2 or FDR (false discovery rate) < 0.05. The Benjamin-Hochberg method was used to correct P-values and to obtain the false discovery rate (FDR). KEGG analysis was conducted on the DEGs identified using BLAST software. To verify the expression of DEGs, the qRT-PCR was to verify the DEGs. Specific quantitative primers were designed using Primer 6.0 software (Supplementary Table 2). The quantitative analysis was performed using Tip Green qPCR SuperMix, and the reaction system contained the upstream and downstream fluorescent quantitative primers (1 µL), TransStart Tip Green qPCR SuperMix (10 µL), cDNA (2 µL), and ddH_2_O (6 µL). The qRT-PCR reaction procedures were shown in Supplementary Table 3. Gene expression levels were determined by the 2^−ΔΔCT^ method, with *Leactin* as the internal reference gene (65).

### Plasmid construction and fungal transformation

The modified pCAMBIA1300-g vector containing the tryptophan decarboxylase (*TDC*) and tryptophan amino transferase genes (*TAA*) was used to construct the RNAi plasmids of *LeTAA* and *LeTDC* genes (66). The promoter of *Legpd* (glyceraldehyde-3-phosphate dehydrogenase) and terminator of T-nos were used to control the expression of *LeTAA* and *LeTDC* and their sense and antisense sequences (65). The RNAi vector was constructed, as described in previous study. The 500 bp antisense fragments on conserved domains of *LeTAA* and *LeTDC* genes were PCR amplified, with the cDNA of S606 as a template. The *Leactin* promoter, antisense fragment, *Legpd* promoter, and the linearized pCAMBIA1300-g plasmid were linked together by homologous recombination to generate *LeTAA-*RNAi (Fig. S1A) and *LeTDC*-RNAi vectors (Fig. S1B) (67). *LeTAA-*RNAi, and *LeTDC*-RNAi were transformed into strains S606 by *A. tumefaciens*-mediated transformation method (67). The empty vector PCAMBIA1300-g was transformed into S606 as control (CK). MYG medium supplemented with 6 μg/mL hygromycin B was used to select positive transformants. Whether the transformants were successfully transformed with *TAA or TDC* fragment was verified by PCR.

### Statistical analysis

SPSS 19.0 (IBM Corporation, United States) was used to determine significant differences between groups and conduct one-way ANOVA and Duncan multiple range tests. p < 0.05 was considered as statistically significant. GraphPad Prism software was employed to plot. Then normal distribution of data was tested by one-sample Kolmogorov-Smirnov test. The data were presented as mean ±SD (standard deviation). Non-normal variables were reported as median (interquartile range [IQR]). Means of 2 continuous normally distributed variables were compared by independent samples Student’ t test. Mann-Whitney U test and Kruskal-Wallis test were employed to compare non-normally-distributed data from multiple groups, respectively.

## Results

### Higher heat tolerance to 40 ℃ heat stress of heat-tolerant strain than heat-sensitive one

In this study, we established a model to compare the heat-tolerance of various *L. edodes* strains. After 40 ℃ heat stress for 0 h, 2 h, 4 h, 6 h, 8 h, 10 h, 12 h and 24 h, the mycelial growth recovery ability of *L. edodes* heat-sensitive strain YS3357 and heat-tolerant strain S606 at 24 ℃ post 7-day culture was investigated (Figure 2). After 10-24 h 40 ℃ heat stress, the strain YS3357 failed to resume growth, while strain S606 resumed growth at all the test time points, indicating that 10 hours of 40 ℃ heat stress is a key point for distinguishing ability of thermotolerance on different *L. edodes* strains.

**Fig. 1.**
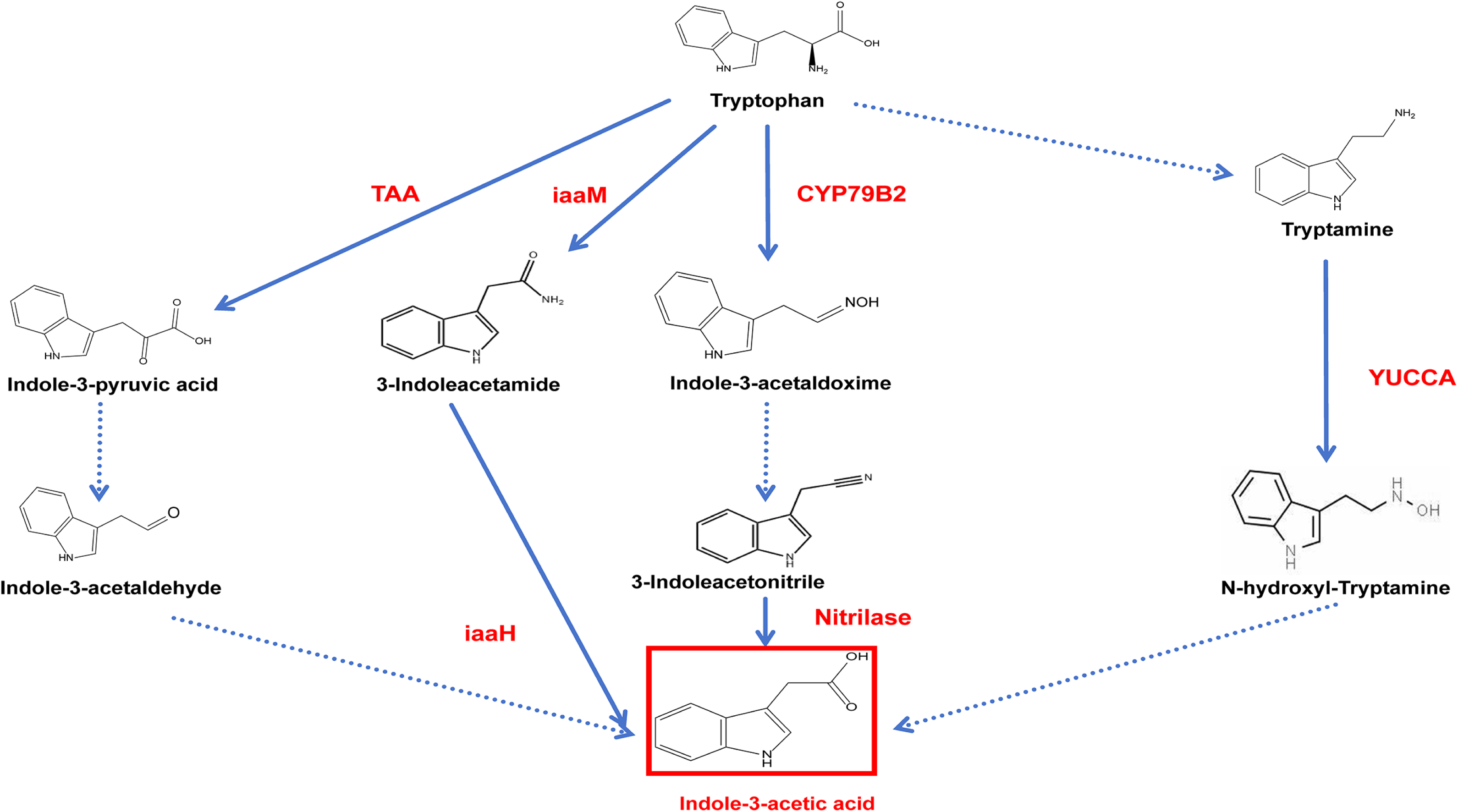
Schematic representation of the auxin biosynthesis pathway in plants. Adapted from Zhao (2010). Key enzymes and intermediates in the auxin synthesis pathway are shown, with relevance to fungal heat tolerance mechanisms.

**Fig. 2.**
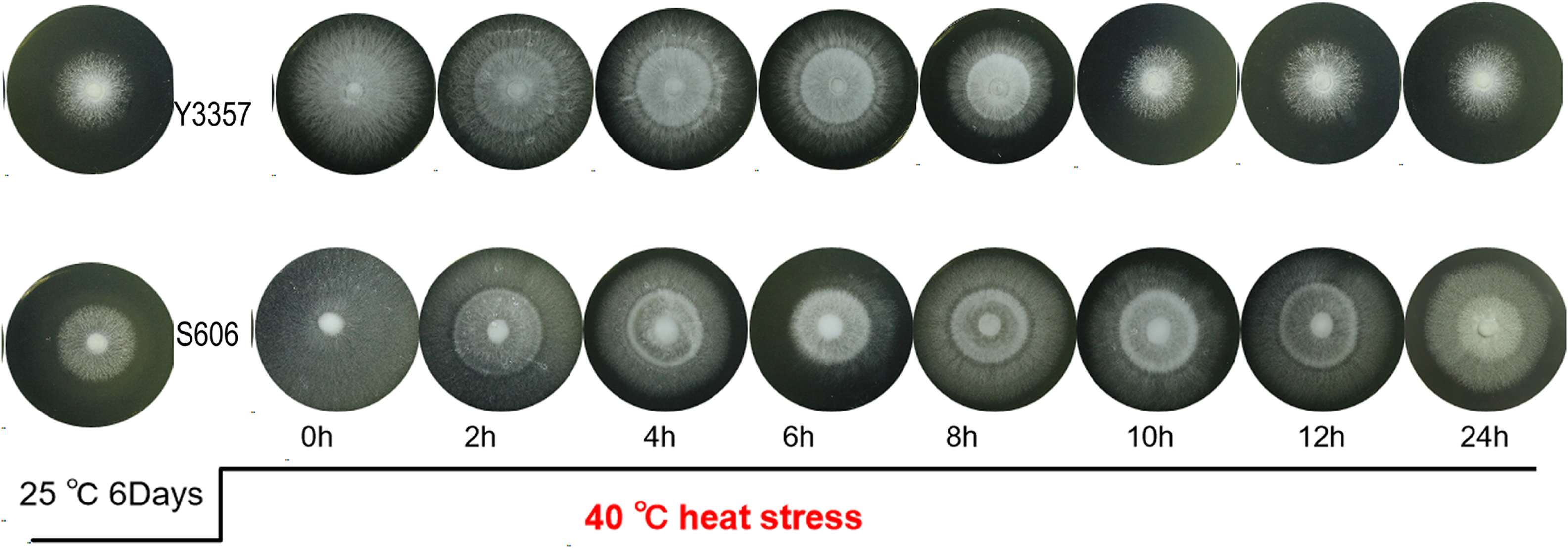
Hyphal growth dynamics under heat stress and recovery. L.edodes strains S606 (heat-tolerant) and Y3357 (heat-sensitive) were cultured at 25 °C for 6 days, subjected to 40 °C heat stress for indicated durations (0–24 h), and allowed to recover for 7 days. Images depict hyphal morphology post-recovery.

### Target metabolomic analysis of phytohormones and related metabolites under different durations of heat stress

To deep understand the correlation between phytohormone and thermotolerance of *L. edodes* strains, we performed the target metabolomic analysis of 9 types of main phytohormones and their related metabolites in *L. edodes* strains S606 and YS3357 hypha after different durations of heat stress. Seven phytohormones including cytokinin (CK), indole-3-acid (IAA), jasmonic acids (JA), gibberellin acids (GA), abscisic acids (ABA), salicylic acid (SA), and ethylene (ETH), as well as 43 phytohormone-related metabolites from *L. edodes* mycelia were successful detected (Supplementary Table 2). Two target phytohormones melatonin (MLT) and(±) Strigolactones (SLs) were undetected from *L. edodes* mycelia.

PCA based on the normalized metabolite expression level and OPLS analysis, the thermotolerance of *L. edodes* strains S606 and YS3357 could be distinguished by the second princIPYAl component (Figure. 3A). The significant changes of metabolite contents were mainly observed after 24-hour 40 ℃ heat stress on two strains, metabolites IAA, Trp, and TAM of both strains showed significant up-regulation under 40℃ of heat stress after 24h, but the increment was greater in heat-tolerant strain S606 than in heat-sensitive strain YS3357, with S606 rising from 2.67 to 113.77ng/g, Y3357 went up from 1.94 to 30.88ng/g, while the contents of SA and JA were increased only in strain YS3357. The analysis of target metabolomics indicated that the heat tolerance of hyphal to 24-h heat stress was highly correlated with the intracellular IAA content, the accumulation of other hormones could not enhance the heat tolerance of *L. edodes* strains (Figure. 3B).

**Fig. 3.**
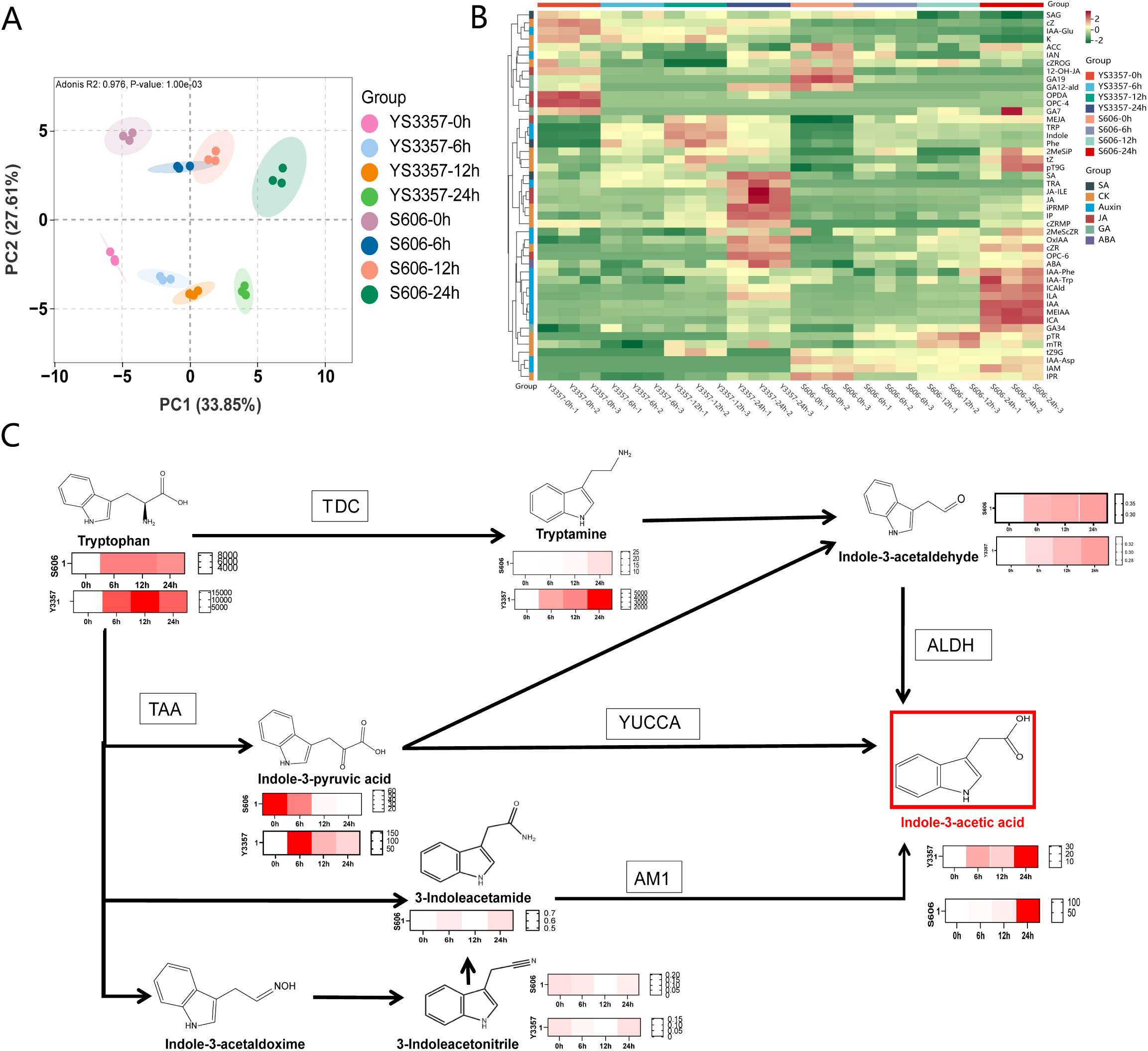
Metabolic profiling of fungal strains during heat stress. (A) PrincIPYAl component analysis (PCA) of differentially accumulated metabolites (DAMs) at 0, 6, 12, and 24 h of 40 °C stress. (B) Heatmap clustering of DAMs across time points. (C) Indole-3-acetic acid (IAA) metabolic flux in S606 versus Y3357 under stress. Color intensity corresponds to metabolite abundance (darker = higher).

### IAA metabolic flux of heat-resistant and heat-sensitive strains was different under heat stress

The biosynthesis pathways of IAA are divided into tryptophan-dependent and tryptophan-independent pathways. According to different intermediate products during IAA synthesis, tryptophan-dependent pathway is further subdivided into IAAOx pathway, TAM pathway, and IPYA pathway in plant, fungi and bacteria. Based on target metabolomics analysis, the intermediates of IAA included IAA, TAM, IPYA, IAN, IAM and IAALD were analyzed. The results showed that the TAM content in Y3357 strain increased from 1426.90 to 6262.77 ng/g after 24h of heat stress, and TAM in S606 increased from 7.4399 to 25.43 ng/g after 24h. The IPYA content of strain S606 was gradually decreased from 61.36 to 12.27 ng/g after heat stress. The IPYA content of strain Y3357 first increased to a peak of 164.1627 ng/g and decreased to 39.79 ng/g after 24h (Supplementary Table 4). The IAALD content increased from 0.26 ng/g to 0.39 ng/g in S606 and from 0.269 ng/g to 0.33 ng/g in Y3357. The IAM and IAN content of both strains were very low before and after heat stress (below 0.1 ng/g). These results indicate that IAA is mainly synthesized through the IPYA and TAM pathway, and the TAM content accumulates rapidly under heat stress, but the increase of IAA content may be mainly synthesized through the IPYA-IAALD pathway (Figure. 3C).

### Silencing of key genes *LeTAA* and *LeTDC* in the IAA synthesis pathway can affect the heat tolerance of *L. edodes* strains

According to the above findings, *L. edodes* strains mainly produce IAA via the IPYA and TAM pathways. To further reveal the IAA synthesis mechanism under heat stress, we investigated the key gene *LeTAA* and *LeTDC* functions in IPYA and TAM pathways. Nine positive transformants each of the *LeTAA* and *LeTDC* genes were obtained based on the two-promoter silencing system, compared with those in wildtype strains S606 and YS3357, the expression levels of *LeTAA* and *LeTDC* genes in transformant-containing strains were decreased by approximately 50%. Three of the transformants (TAA-RNAi-Y3357-12, TAA-RNAi-S606-13 and TDC-RNAi-S606-9) were selected with wild-type S606 for subsequent analyses. The results showed that there were no significant differences in mycelial growth rate and colony morphology on MYG medium between 3 transformant strains and 2 wildtype strains (S606) (p > 0.05) (Figure. 4A). These results suggested that the interference with *LeTAA* and *LeTDC* genes had no significant influence on *L. edodes* strain growth. At 24 h post 40 ℃ heat stress, TAA-RNAi-S606-13 and TDC-RNAi-S606-9 transformant strains could not resume growth, at 12 h, transformant strains resumed growth, but their recovery phenotype was inferior to that of the wildtype S606, indicating the interference of *LeTAA* and *LeTDC* genes in heat-tolerant strains diminished their heat resistance (Figure. 4B-C). Since it mainly synthesized IAA through the TAM and IPYA pathways, the interference of the genes of TAA and TDC might have attenuated the heat tolerance of the mushroom by affecting the synthesis of IAA (Figure. 4D).

**Fig. 4.**
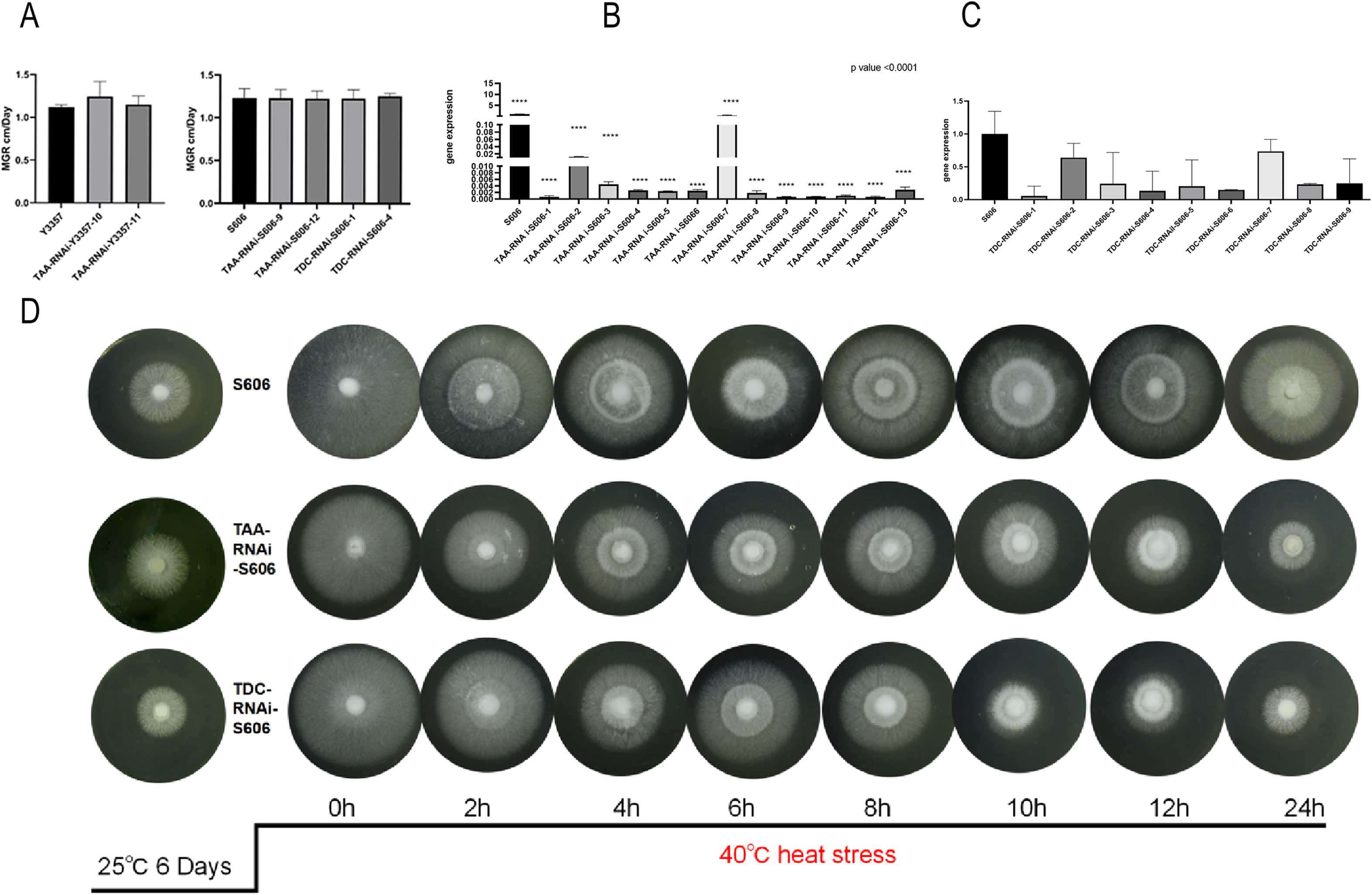
Transcriptomic analysis of heat stress responses. (A) PCA of global gene expression and Venn diagram of strain-specific genes. (B) Differentially expressed genes (DEGs) at each stress time point. (C) Expression trends of heat-responsive genes and enriched KEGG pathways (FDR < 0.05).

### Analysis of gene expression changes during 0-24h of 40℃ heat stress in *L. edodes* **strains**

We further performed a transcriptomic analysis of the expression of the two strain genes during 0-24h (0, 2, 4, 6, 12, 24h of heat stress at 40 ℃). PCA analysis showed that the gene expression patterns of S606 and YS3357 strains were clearly separated in the second component, and then in the first component after heat stress (Figure. 5A). It can be seen in the Upset plot that the number changes of DEGs in the two strains showed different patterns at different times of heat stress, with the heat-resistant strain S606 increasing with increasing heat stress time, peaking of 4582 genes at 24 h. The heat-sensitive strain Y3357 produced a high number 4111 of differential genes after 2h of heat stress (1.5 times that of S606), At a peak of 5817 genes after 24h of heat shock, indicating that gene expression is unstable under YS3357 heat stress (Figure. 5D-E). In heat-tolerant strain S606, 1486 genes were up-regulated after 2h of heat stress (cluster 3, 4), which were mainly related to protein-related mitochondrial localization and target to the mitochondria, protein folding, protein refolding, cell response to heat, temperature stimulation pathways. However, the gene expression pattern of heat-sensitive strain Y3357 in response to heat stress was quite different from that of S606. About 50% of the genes were significantly down regulated after 2h of heat stress, and only 792 genes (cluster 5) were up-regulated after 2h of heat stress, and the up-regulated genes were mainly related to protein folding and refolding pathways. These results indicate that the heat-resistant strain S606 gained thermostability by the earlier up-regulated gene expression strategy (Figure. 5F-G).

**Fig. 5.**
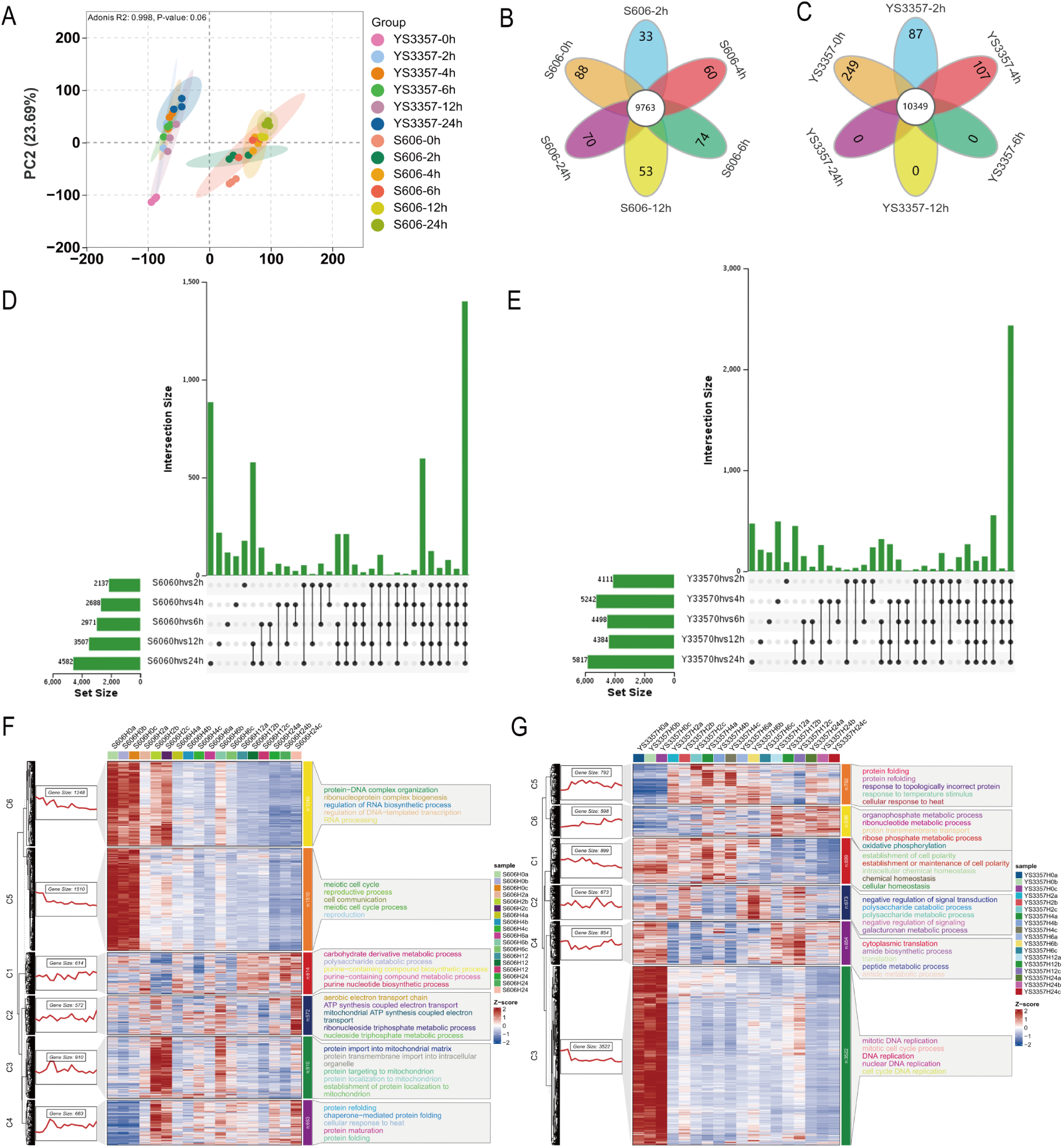
Functional validation of heat tolerance-associated genes. (A) Growth rates of S606, and transformants (TAA-RNAi and TDC-RNAi). (B, C) Relative expression of LeTAA and LeTDC in S606 versus transformants (***p < 0.001, t-test). (D) Recovery phenotypes after 40 °C shock. Data represent mean ± SD (n = 6).

### *L. edodes* strains exhibited enhanced heat tolerance through IAA-mediated regulation of the MAPK signal transduction pathway during 40 **°C** heat stress 0-24h

To investigate the mechanisms underlying the remarkable heat tolerance of the S606 strain during heat stress, this study employed a nine-quadrant analysis to compare the differential expressed genes (DEGs) (Supplementary Table 5, 6) between S606 and Y3357 at 0-24 hours post-heat exposure (Figure.7 A-C). The analysis indicated that genes situated in the first and second quadrants, which were highly expressed in S606 but either down-regulated or unchanged in Y3357 following heat stress, were closely associated with the superior heat tolerance observed in S606. Notably, after 6, 12, and 24 hours of heat shock, the number of genes in these quadrants increased, suggesting that the S606 strain effectively mitigated heat stress by up-regulating a greater number of genes compared to the Y3357 strain. KEGG enrichment analysis of gene functions in the first and second quadrants demonstrated distinct functional changes in highly DEGs in S606 strain during 0 -24h of heat stress. KEGG enrichment analysis of gene functions in the first and second quadrants revealed distinct functional changes in highly DEGs in S606 strains during 0-24h heat stress. In terms of metabolic pathways, highly DEGs of S606 were mainly concentrated in amino acids and secondary metabolites after 6h of heat stress. By 12h of heat stress, DEGs in S606 strains were distributed in the tryptophan metabolism pathway, alcohol, and lactic acid metabolism, among which *ALDH* genes in the IAA synthesis pathway also showed high expression differences. This was consistent with the increase of auxin IAA content thereafter. In the signaling pathways, after 6 hours of heat stress, the *Hog 1* gene in the hyperpermeability pathway and the *MKK 12* gene in the cell wall response pathway of S606 strain were highly expressed. By 12 hours of heat stress, the *FUS 3* gene in the pheromone pathway, the WSC123 receptor in the cell wall response pathway, *Pbs 2*, *Hog 1*, *Gpd 1*, *Gre 2* in the hyperpermeable pathway, and various genes in the starvation pathway were highly expressed. After 24 hours of heat stress, fourteen genes in high permeability pathways, cell wall response pathway, pheromone receptor pathway, and starvation MAPK signaling pathway, as well as DNA mismatch and recombinant repair, showed high differential expression. These results suggest that the heat-tolerant strain S606 improves heat tolerance by regulating secondary metabolic pathways and signaling pathways responding to stress, as well as the expression of genes related to DNA repair. The simultaneous analysis of promoter regulatory elements in the upstream regions of signal regulation-related genes in S606 identified auxin response elements in the promoters of *WSC123*, *Ste 3*, *Cdc 24*, *Cla 4*, *Rom 1, 2*, *FUS 3*, *Pbs 2*, *Hog 1*, and the cell growth cycle transcription factors *Tec 1*, *Ste 12*, *clb 1/2*, and *Gre 2*. This indicates that the S606 strain improved heat tolerance by increasing the expression of genes related to IAA content regulation and stress response and MAPK signaling response pathway at 24h after heat stress (Figure. 6).

**Fig. 6.**
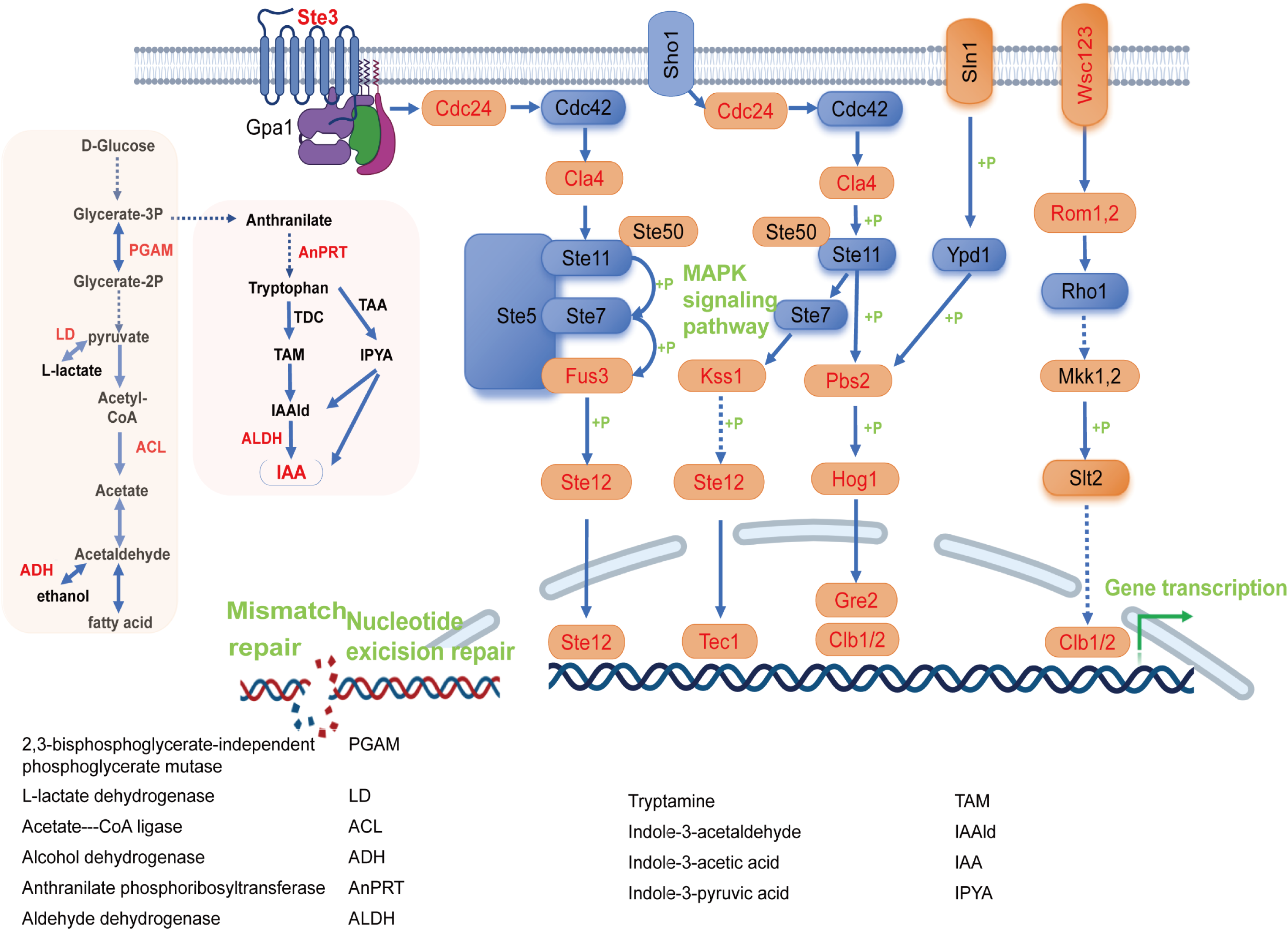
Proposed model of IAA-mediated MAPK pathway regulation under heat stress. Red-labeled pathways (fatty acid metabolism, IAA synthesis) indicate upregulated genes. ARF-responsive elements (red) in MAPK components suggest auxin crosstalk. Gene acronyms: ARF (Auxin Response Factor), MAPK (Mitogen-Activated Protein Kinase), TAA (Tryptophan Aminotransferase), TDC (Tryptophan Decarboxylase).

**Fig. 7.**
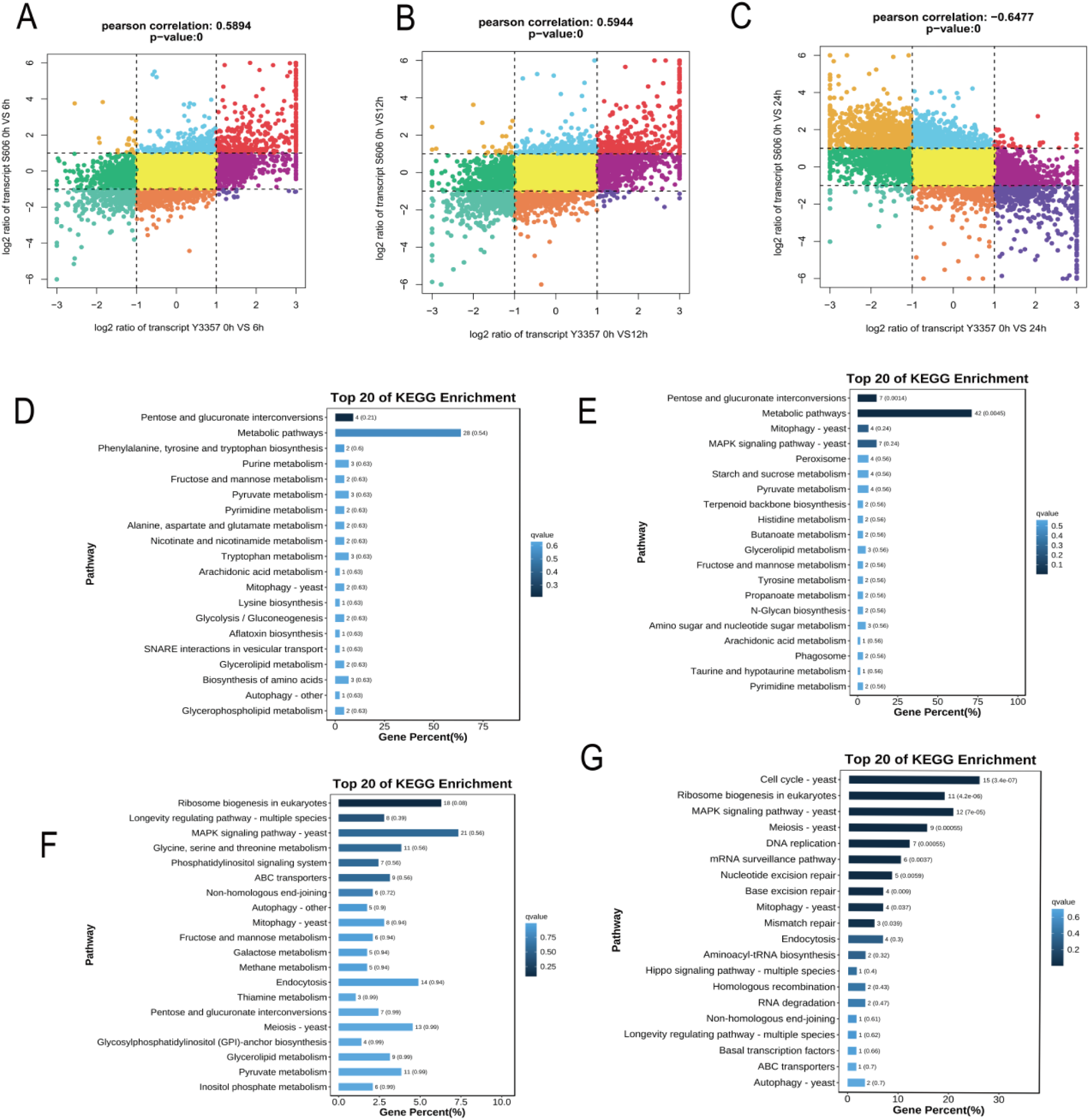
(A) Growth rates between Y3357 and S606 and their transformants. (B)Comparison of IAA content between Y3357, S606, and their transformants. (C)Recovery growth phenotypes of S606 and its transformants after heat shock at different times over 6 days. (E)comparison of gene expression levels between Y3357 and its transformants. (D)Recovery growth phenotypes of Y3357 and transformants after heat shock at different times over 6 days. (F)comparison of gene expression levels between Y3357 and its transformants.

### Association analysis of the expression levels of transcriptome and hormone metabolites

To explore the effect of gene expression and the expression level of hormone metabolites on heat tolerance after heat stress, we performed OP2LS analysis. KEGG enrichment analysis of genes and metabolites obtained from OP2LS analysis revealed that genes related to metabolism during heat stress in S606 were significantly enriched in repair pathways, including cell cycle, cell autophagy, and homologous recombination (Figure.7 D-E). Notably, the expression of energy-related intermediate metabolite synthesis pathways, such as acetyl-CoA, increased significantly. These findings suggest that S606 improved heat tolerance during heat stress by enhancing cell cycle, autophagy, and homologous recombination, as well as energy metabolism and other pathways. The results of the OP2LS analysis are highly consistent with those of the nine-quadrant analysis, which indicated that the S606 strain gained stronger heat tolerance by positively regulating the expression of stress signaling pathway, DNA repair, and energy metabolism-related genes. Furthermore, the increase in auxin IAA content supports the notion that S606 improved heat tolerance by modulating stress response and energy metabolism pathways.

## Discussion

As mentioned in the literature review, IAA synthesis in plants and bacteria is mainly metabolized through the Trp-dependent TAM, IPYA, IAN, IAM pathways. Several reports have shown that yeast *DMKU-CP293* strain produced IPYA as the main pathway, and IAM and TAM as the complementary pathway (68). The current study found that *Rhizobium tropici* and *Fusarium species* synthesize IAA via the IPYA, TAM, and IAM pathways. Additionally, the IAM pathway has been identified as a principal route for IAA synthesis in anthracis (*Colletotrichum gloeosporioides* (69), *C. acutatum* (70), *C. fructicola* (71)). In *Magnaporthe oryzae*, IAA is mainly synthesized through the IPYA pathway (72). Prior studies that noted that fungal intracellular IAA biosynthesis predominantly proceeds through three Trp-dependent pathways: TAM, IPYA, and IAM.

In this study, we assessed the levels of tryptophan along with metabolites associated with the TAM, IPYA, and IAM pathways at various time points (0h, 6h,12h, 24h) during heat stress at 40 °C. What is surprising is that the concentrations of IAN and IAM pathways remained low and unchanged throughout this period. Despite identifying genes related to both the IAN and IAM pathways within mushroom genomes, metabolite detection revealed that these pathways were not efficient in synthesizing IAA. As far as we aware this is the first time in *L. edodes* strains that Trp-dependent synthesis of IAA, which primarily occurred via two key routes: TAM and IPYA pathways.

Both previously published data alongside quantitative metabolomics analyses from this experiment demonstrated a significant increase in intracellular IAA content in *L. edodes* between 12 to 24 hours post-heat stress exposure, indicating an enhancement in IAA biosynthesis under prolonged thermal conditions. Targeted metabolomics assessments measuring precursor levels for IAA synthesis in S606 strains subjected to heat stress at 40 °C over intervals of 0h, 6h, 12h, and 24h revealed substantial alterations in metabolites linked to the IAA synthesis pathway under heat exposure. Notably among these changes was a marked increase in levels of Trp, TAM, and IAALD after six hours of heat stress, conversely, the concentration of IPYA exhibited a decline. The elevated IAA content was observed after 12 hours of heat stress, with the timing of these changes significantly lagging behind that of anabolic intermediates. The most striking result is that the tryptophan content in the YS3357 strain is 175 to 240 times higher than that in S606. This abnormal accumulation of tryptophan may be attributed to its inability to synthesize other metabolites and its ineffective conversion into IAA. This observation may support the hypothesis that high intracellular concentration of TAM can exert a toxic effect on cells, which may represent a critical factor distinguishing YS3357 from S606. However, further experimental evidence is required to determine whether this is a common occurrence among heat-sensitive strains.

Tryptophan transferase (*TAA)* facilitates the conversion of Trp into IPYA, while tryptophan decarboxylase (*TDC*) converts it into TAM, both are essential genes regulating IAA synthesis in various organisms. Previous studies have reported that *TAA* gene regulates IPYA synthesis in *Ustilago maydis* strain (53) and *Leptosphaeria maculans* (54) strain, the *Umtam1* and *tam2* deletion mutants (the tryptophan aminotransferase genes) exhibit the decreased IPYA content, and exogenous addition of Trp rescues the IPYA content decrease, the expression of gene *LmTAM1* was upregulated after the addition of tryptophan, its expression was positively correlated with IAA production. *Taphrina deformans* (73), *Metarhizium robertsii* (74), and *Cyanodermella asteris* (75)strains have been reported to possess *TDC* activity, and the exogenous addition of TAM in *Azospirillium*, *Ustilago maydis* (73), and *Leptosphaeria maculans* strains (54) enhances *TDC* activity, thus increasing the level of IAA. Overall, research on *TAA* and *TDC* genes within fungal species remains limited. In this study, we silenced the expression of both *LeTAA* and *LeTDC* genes in the *L. edodes* S606 strain. Our findings indicate that downregulation of these genes does not adversely affect normal hyphal growth but does reduce heat resistance, suggesting that the expression levels of *LeTAA* and *LeTDC* play a significant role in modulating hyphal thermotolerance.

Furthermore, fungal MAPK signaling pathway encompasses pheromone signaling, cell wall integrity signaling, as well as responses to salt and hypertonic stress conditions. The fungal cell wall signaling cascade and hypertonic pathways typically respond to compensatory mechanisms associated with heat stress, resulting in the accumulation of intracellular trehalose (76-78). This process enhances intracellular osmotic pressure and leads to plasma membrane stretching (79-81). WSC receptors in yeast are capable of responding to heat stress and rapidly reacting to cell wall stress signals (82, 83), they also influence the hyphal growth and heat sensitivity (84). In this study, we performed a nine-quadrant analysis at 2, 4, 6, 12, and 24 hours post-heat stress. The remarkable results revealed that most genes involved in the MAPK signal transduction pathway, as well as those related to lactate, alcohol, and IAA synthesis in the *L.edodes* strain S606 was significantly expressed after 12–24 hours of heat exposure. In contrast, these gene expression levels were either down-regulated or remained unchanged in the heat-sensitive strain YS3357. Within the MAPK signal transduction pathway, we identified ARF response elements within the promoter regions of thirteen genes across various pathways (see attachment). It may be the case therefore that during the period of 12–24 hours under heat stress conditions, auxin IAA positively regulates the MAPK signal transduction pathway to initiate the expression of stress-related genes (Supplementary Table 7).

In conclusion, this paper investigates alterations in auxin synthesis pathways and related to auxin synthesis metabolite changes under conditions of heat stress through transcriptomics approaches at different time points during thermal exposure. Our findings would seem to show that intracellular IAA synthesis primarily occurs via both TAM and IPYA pathways in *L. edodes*, however, excessive TAM accumulation after heat stress contributes to decreased thermotolerance. Furthermore, increased levels of IAA are predominantly derived from IPYA-synthesized IAALD. Gene silencing experiments targeting *LeTAA* and *LeTDC* within the auxin synthesis pathway demonstrated reduced thermotolerance in hyphae. Notably, elevated intracellular IAA content observed between 12–24 hours during heat stress positively regulated genes implicated in the MAPK signal transduction pathway as a protective response against thermal damage. These results provide a deeper understanding of the mechanism of IAA synthesize in *L. edodes* strains, providing a molecular basis for future breeding of thermotolerant strains.

## Acknowledgements

This work was financially supported by the National Key R&D Program of China (Grant No. 2023YFF1000804), and the National Natural Science Foundation of China (Grant No. 32072641)

## FIGURE LEGENDS

Fig. S1. Agrobacterium transformation vectors. (A) TAA-RNAi vector; (B) TDC-RNAi vector.

Fig. S2. LeTAA segments in Y3357 and S606 strains. M: BM 1kb DNA marker; Lanes 1-2: LeTAA amplification from Y3357; Lanes 4-5: LeTAA amplification from S606.

Fig. S3. LeTDC fragments in Y3357 and S606 strains. M1: BM 1kb DNA marker; M2: BM2000+ DNA marker; Lanes 1-3: LeTDC amplification from Y3357; Lanes 4-6: LeTDC amplification from S606.

## TABLE LEGENDS

Table S1 UPLC-MS/MS instrumentation details. System configuration: ExionLC™ AD UPLC coupled to QTRAP® 6500+ MS/MS. Table S2 qRT-PCR primers. Primer sequences for LeTAA, LeTDC, and reference genes (e.g., actin). Table S3 qRT-PCR thermal cycling conditions. Protocol: Initial denaturation at 95 °C (3 min), 40 cycles of 95 °C (10 s), 60 °C (30 s). Table S4 Quantification of plant hormones. Concentrations of IAA and related metabolites measured via UPLC-MS/MS. Data are mean ± SD (n = 3).

Table S5 Enriched DEGs in S606 clusters. KEGG pathways and GO terms enriched in heat-tolerant strain S606 (FDR < 0.05). Table S6 Enriched DEGs in Y3357 clusters. KEGG pathways and GO terms enriched in heat-sensitive strain Y3357 (FDR < 0.05). Table S7 Comparative analysis of MAPK pathway gene number and activation elements in heat-resistant vs. heat-sensitive strains post-heat stress

## Data availability

The raw transcriptome sequencing data generated in this study have been deposited in the Genome Sequence Archive (GSA) at the National Genomics Data Center (China National Center for Bioinformation) under BioProject accession number CRA016083 (Submission ID: subCRA025741). The dataset is publicly accessible via: https://ngdc.cncb.ac.cn/gsa.

Submission date: April 22, 2024; Status: Released (Checked OK).

